# Bio-Mol:Pretraining Multimodality Bioactivity Profile for Enhancing Small Molecule Property Prediction

**DOI:** 10.1101/2023.11.02.565401

**Authors:** Hiu Fung Yip, Xiao Wei, Zeming Li, Qing Ren, DongSheng Cao, Lu Zhang, Aiping Lu

## Abstract

Non-optimized pharmacokinetic parameters serve as the primary cause of failure in clinical trials of drugs. Therefore, the successful prediction of pharmacokinetic parameters during the pre-clinical stage is crucial for the success of drug candidates. Conventional methods primarily rely on 2D structural information, while advanced models extend the features to other structural-related information or use advanced computational models to improve prediction accuracy. However, to gain a comprehensive understanding of small molecules, integrating bioactivity profiles with chemical structural information is essential. One significant challenge in this integration is the high proportion of missing values within experimentally validated bioactivity profiles for most small molecules. To address this challenge, we introduce Bio-Mol, an artificial intelligence model designed to effectively handle this issue. Bio-Mol utilizes a pretrain and finetune strategy, enabling the incorporation of a large proportion of missing bioactivity profiles during the small molecule representation learning process.

Comprehensive evaluations of Bio-Mol demonstrate a notable improvement in predicting molecule properties. The integration of missing bioactivity profiles enhances the AUROC of average 5.2% compared to the previous state-of-the-art model’s predictions. Furthermore, we explore the potential of Bio-Mol in predicting synergistic drug combinations, highlighting its versatility and broader applications in the field of drug discovery.

The successful implementation of Bio-Mol showcases its efficacy in overcoming the challenges posed by missing bioactivity profile data. This model paves the way for optimizing small molecule pharmacokinetics prediction, providing valuable insights for drug development and discovery processes.

## 1 Introduction

Small molecule drug discovery is crucial for improving human health, but it is time-consuming and has a high failure rate. 90% of drug candidates would fail after entering clinical studies with unpredictable experimental factors [4, 16]. The main reasons for the failure are 1) lack of clinical efficacy (40%-50%), 2) unmanageable toxicity (30%), 3) poor drug-like properties (10%-15%) [4, 9]. Those failures could be largely explained by issues rooted in early discovery such as inadequate target validation or suboptimal ligand properties and non-optimized pharmacokinetics properties prediction [15]. Hence, optimizing the small molecule drug pharmacokinetics properties, namely, Absorption, Distribution, Metabolism, Excretion, and Toxicity (ADMET) is the key to accelerating the small molecule drug discovery process.

Computational-Aid Drug Discovery (CADD) has demonstrated its outstanding performance in small molecule ADMET prediction. The reasons for the improved performance are the following; First, structural information of small molecules could predict the ADMET. Second, similar chemical structures would share similar pharmacokinetics properties, which is usually called the “similarity principle”. A landmark study has demonstrated the potential of CADD in ADMET prediction, for example, Fang, X. et al. proposed a graph neural network to refine the molecule representation by small molecule’s 2D geometric information [6].

However, Bioactivity profiles are continually proven to provide complementary information to the structural information of a small molecule. Haghighi, M. et al. showed that gene expression and cell morphology features provided complementary information to improve the prediction of the mechanism of action (MoA) of a drug [8]. In this study, we will integrate the most comprehensive bioactivity profiles for small molecules. A small molecule could be represented by five layers, namely, A) chemistry, B) targets, C) networks, D) cells, and E) clinics, each layer could be further classified into five sub-layers, and each sub-layer is represented by a 128-dimensional vector, the definition is strictly follow Duran-Frigola, M. et al. Work [5].To the best of our knowledge, They are the first to show the “similarity principle” is also applicable to the bioactivity profile of small molecules.

Although the Bioactivity profiles could provide a full spectrum of features of the small molecule, which could improve the ADMET prediction, only a small fraction of studies include that information in the ADMET prediction model. The main reason is the bioactivity profiles are missing for most of the small molecules (up to 99.9%). Such high proportions of missing values in the bioactivity profiles make them not suitable for developing a general small molecule ADMET prediction model which targets to virtual screening for potential clinical trial candidates from a massive amount of chemical space.

The existing method that considers the bioactivity profile in the small molecules ADMET prediction is to impute the miss values in the small molecule bioactivity profile, the landmark study of this bioactivity profile imputation is Bertoni, M. [1], they proposed a Siamese Neural Network (SNN) model to impute the missing values for the 1 million small molecules curated from the database, and demonstrated the improved AUROC in the 9 benchmark ADMET prediction tasks compare to the classical structural based methods.

However, there are two major drawbacks in the SNN imputation, 1) imputation could cause error accumulation; statistical guidance articles have stated that if more than 40% data are missing in important variables, the results should only be considered as hypothesis-generating [3, 11]. 2) SNN is not an end-to-end model; it does not include the molecular property prediction tasks to refine their imputation result, but molecules with similar properties should have a closer distance in the imputed profile to those that are not.

Here, we develop Bio-Mol: a multi-modality contrastive learning model for pretraining the bioactivity profile of small molecule representation. This model is to address the major challenge of integrating the high proportion of missing values of bioactivity profile in small molecules to improve the ADMET prediction. The key ideas are 1) Different modalities have strong relationships, so the masked modality could be reconstructed through the non-masked modality. 2) a small molecule with two modality masks should be closer than two with the same modality mask. 3) The downstream task should be used to finetune the pretrained latent space.

To address the challenge, we employed a pertain and fine-tune strategy. In the pretraining stage, we trained on 1 million small molecules lacking labeled ADMET properties to learn low-dimensional representations for each molecule. We utilized masked input imputation, simulating missing proportions, and trained the model to impute the masked sub-layers. Additionally, a triplet loss was deployed to encourage molecules with different masks to have closer distances in the latent space. In the finetune stage, we refined the pretrained representation with downstream ADMET tasks, employing cross-entropy loss. This ensured that molecules with similar ADMET properties shared similar low-dimensional representations, enhancing prediction accuracy.

Our Bio-Mol significantly improves the AUROC compared to SNN, and the other structure-based baselines molecule property prediction methods. Moreover, our model can impute the masked modalities with low MSE. Our model is open source and stored in GitHub: https://github.com/YIPhiufung1997/Bio-Mol.

## 2 Method

### 2.1 Pretraining Dataset

In the pretraining phase, we pretrain Bio-Mol using 1,000,000 unlabelled molecules sampled from Chemical Checker [5], a public access database containing small molecules five layers of information, namely, A) chemistry, B) targets, C) networks, D) cells and E) clinics, each layer could further classify into five sub-layers, each sub-layer is represented by a 128-dimensional vector. We select the processed data without imputation, so there are missing values for each sub-layer and each small molecule.

### 2.2 Contrastive-based pretraining

The pretraining stage is the implementation of the two ideas: 1) different modalities have strong relationships, so the masked modality could be reconstructed through the non-masked modality. 2) a small molecule with two modality masks should be closer than two with the same modality mask.

We utilize high missing value chemo-bioactivity profile to pretrain Bio-Mol. For each small molecule, we got all its non-missing value sub-layers. Then, we generate two independent masks to mask the sub-layers according to the observed missing proportion of the sub-layer. Denote M as the masked function,

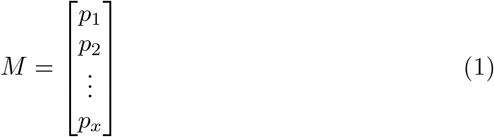

*p*_*x*_ is the observed missing probability of sub-layer *x*, where *x* = 1, 2, …25, 0 *≤ p*_*x*_ *≤* 1. And for each small molecule, we will generate the following two masks:

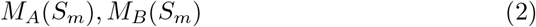

where *S*_*m*_ is the small molecule *m, m* = 1, 2, 3, …, *N* . *N* is the total number of small molecules in the pretrain stage. *M*_*A*_, *M*_*B*_ are the two independent masks to mask the sub-layers according to the observed missing proportion of the sub-layer. Similarly, we can define the different small molecules with the same mask as:

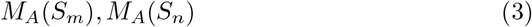

where *S*_*m*_, *S*_*n*_ are the two small molecules with the same mask, where *m* = *n* and *m, n* = 1, 2, …, *N* . In order to let *m* and *n* be structurally similar, we perform KNN search to ensure, the small molecule *n* is the K nearest neighbor of the small molecule *m* in the sub-layer A1 (2D structure).

### 2.3 AutoEncoder architecture

Encoder: Our encoder is composed of eight linear fully connected layers; each layer is followed with a ReLU activation function. Denote as a function *E*(*x*) where *x* is the input. Decoder: Our encoder is composed of five linear fully connected layers; each layer is followed with a ReLU activation function. Denote as a function *D*(*x*) where *x* is the input. For each small molecule *S*_*m*_, we will denote the final output of the decoder as Ŝ_*m*_,

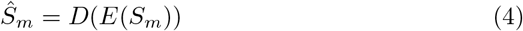

Now, we can define the loss function in the pretrain stage as,

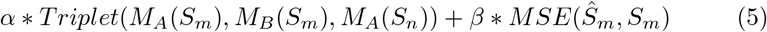

where *α, β* are hyperparameters, *Triplet*(*M*_*A*_(*S*_*m*_), *M*_*A*_(*S*_*m*_, *M*_*A*_(*S*_*n*_)) represents the triplet loss that the distance of same molecule with different masks, and maximize the distance of two molecules with same masks. *MSE*(Ŝ_*m*_, *S*_*m*_) is the mean square error being to reconstruct the original input data by filling in the missing values or imputing the non-masked modalities based on the information learned from the masked modalities.

### 2.4 ADMET tasks finetuning

We use the benchmark datasets provided in MoleculeNet [17], including both classification and regression tasks. To test our model ability on out-of-distribution samples, we follow the Scaffold split strategy as recommended in [2]. Scaffold split is a chemical structure-specific splitting strategy which considered to be more challenging than random split, and it could better simulate the data distribution during the drug discovery process. In this work, the Train/validation/test proportion is 8:1:1 among all baselines.

### 2.5 Evaluation experimental setup

We compare Bio-Mol to SNN [1], classical structural-based methods, and other competitors. Here, we compare the AUROC of the baseline methods in 4 ADMET classification tasks. The computations were executed on a server with the following specifications: AMD EPYC 7302 (2S/16C) @3.0GHz/1T RAM, 2 sets of Nvidia Ampere A100 40G, Oracle Linux 8.4 and CUDA 11.4.

### 2.6 Baselines selection

We compare the proposed method with various competitive baselines. SNN, GROVER, GEM, GraphMVP, Classic. SNN and Classic are run by us. GROVER, GEM, and GraphMVP results are based on the [7].

### 2.7 Evaluation metric

During the evaluation phase, we calculate the AUROC for each classification dataset and then take the average across all datasets. This approach provides a holistic assessment of the models’ performance, considering their effectiveness across various classification tasks within the MoleculeNet dataset.

## 3 Result

### 3.1 Advancing Small Molecule Analysis: Embracing the Full Spectrum of Features

In this study, we developed a method that incorporates diverse features, including structure, target, cell response, network, and clinical data, to gain a holistic understanding of drug interactions (Fig. 1A upper panel). Our approach goes beyond existing methods by considering the whole spectrum of small molecule features, leading to improved predictions and enhanced therapeutic strategies.

**Fig. 1.**
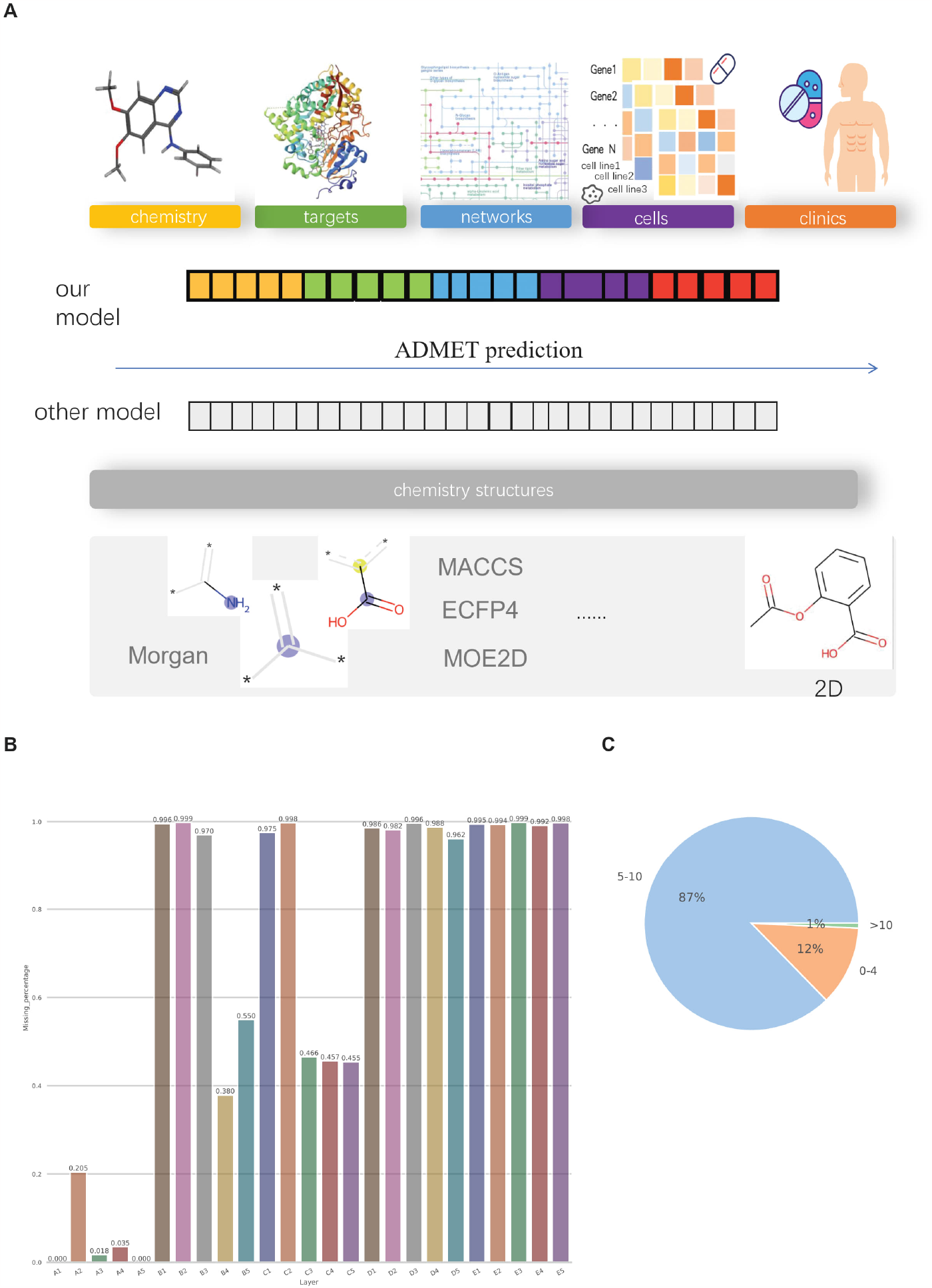
Unique small molecule representation with bioactivity profile and the missing percentage of the bioactivity profile. A) Distinguishing small molecule representation with bioactivity profile (Upper panel) from structure-only models (Lower panel). B) The bar plot displays the percentage of small molecules that lack information regarding a specific sub-layer (A1-E5); The height of each bar represents the proportion of small molecules without sub-layer information. C) The pie chart illustrates the distribution of small molecules based on the number of sub-layers; “0-4”: Represents the percentage of small molecules with 0 to 4 sub-layers.”5-10”: Represents the percentage of small molecules with 5 to 10 sub-layers.” 10”: Represents the percentage of small molecules with more than 10 sub-layers.

An example similar to our method is the Structured Neural Network (SNN), which also incorporates multiple data types. However, SNN primarily focuses on imputation, which can be prone to errors. On the other hand, structure-based methods, such as GROVER, GEM, and GraphMVP rely on analyzing molecular structures (Fig. 1A Lower panel) using graph-based representations and similarity measures. However, they overlook other relevant factors, such as target interactions and other molecular properties.

### 3.2 Integrating Unexplored Chemo-Bioactivity Features

Due to the substantial amount of missing information and the potential introduction of bias, other methods do not incorporate target, cell response, network, and clinical data into their models. As shown in Fig. 1B, the missing percentage for these features ranges from 1.8% (A3) to 99% (E3). Moreover, an analysis of small molecules reveals that 87% of them possess only five to ten sub-layers of information, while 12% have 0-4 sub-layers, and just 1% have over 10 sub-layers (Fig. 1C). The significant presence of missing values and the limited availability of comprehensive data poses a significant challenge. Including these features directly in the model without accounting for the missing data can introduce biases and inaccuracies in the predictions. Consequently, most methods choose to exclude these features to ensure reliable results and avoid potential distortions.

In contrast to other methods, we recognize the significance of incorporating the bioactivity profile into molecule property prediction tasks. To address the challenge posed by the high percentage of missing information, we employ a pretrain and finetune strategy to encode this valuable data into our model (Fig. 2).

**Fig. 2.**
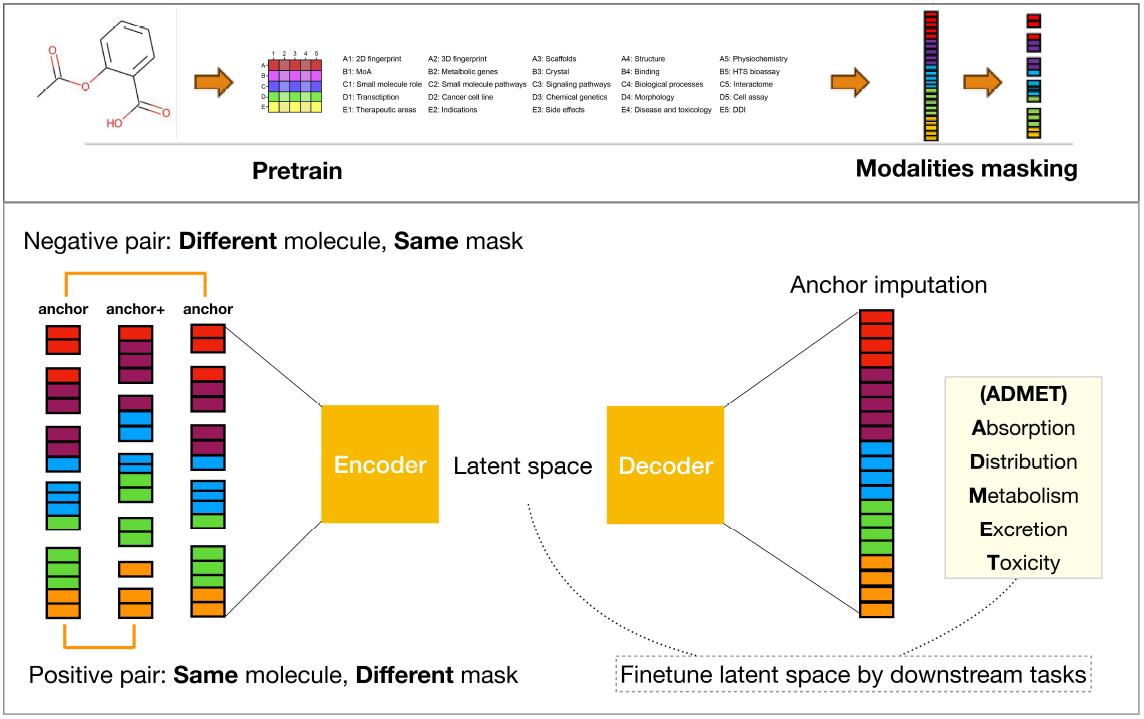
Pretrain-finetuning strategy for small molecule bioactivity profile.

By leveraging this strategy, we can effectively capture and utilize the bioactivity profile despite the presence of missing values. The pretraining phase allows the model to learn from a large dataset, including molecules with varying degrees of missing information. This process enables the model to capture patterns and relationships within the data, even when certain features are incomplete.

### 3.3 Our method improves the performance of molecule property prediction

The overall performance of our method and the other baseline methods are summarized in (Table 1). We have the following observations: (1) Bio-Mol demonstrated the highest AUROC in on all 3 out of 4 datasets (Table 1). On the classification tasks, Bio-Mol achieves an overall relative improvement of 5.2% on the average ROC-AUC compared with the GEM. (2) Clinical Toxicity is the only task GEM outperforms our method, which is 0.1% higher than our method. The highest AUROC task is BACE (mean=0.955, sd=0.015), which includes the experimentally tested for their inhibitory activity against the beta-secretase enzyme. The Lowest AUROC task is Sider, which predicts the side effect record from the experimental validation (mean=0.684, sd=0.008).

**Table 1.** Overall performance of different methods for classification tasks.

| Structure + Bioactivity* |  |  |  |  |
| --- | --- | --- | --- | --- |
| Metric: AUROC |  |  |  |  |
| Splitting method: Scaffold split*** |  |  |  |  |
| Method/Task | Clinical Toxicity | BBBP | BACE | SIDER |
| No. samples | 1,478 | 2,039 | 1,513 | 1,427 |
| our model | 0.902(0.016) | **0.951(0.006)** | **0.955(0.015)** | **0.684(0.008)** |
| SNN | 0.602(0.031) | 0.851(0.025) | 0.732(0.018) | 0.550(0.010) |
| Classic | 0.565(0.035) | 0.610(0.012) | 0.560(0.003) | 0.537(0.042) |
| GROVER | 0.703(0.137) | 0.868(0.022) | 0.824(0.036) | 0.612(0.025) |
| GEM | **0.903(0.007)** | 0.888(0.004) | 0.879(0.011) | 0.632(0.015) |
| GraphMVP | 0.791(0.028) | 0.724(0.016) | 0.812(0.009) | 0.639(0.012) |

## 4 Discussion

In this study, we aimed to investigate the potential of bio-fingerprints in enhancing the prediction performance of Absorption, Distribution, Metabolism, and Excretion (ADME) as well as toxicity. To validate our hypothesis, we conducted a comprehensive comparison with various baseline methods targeting different objectives. Firstly, we compared our method against the classical molecular fingerprint-based method called Classic [1], and our approach outperformed it. Secondly, we evaluated our method against a structural-based method GROVER [14], GEM [6] and GraphMVP [12]. Also, we compared our model with other models with bioactivity profiles, and SNN baseline [5]. Our model consistently achieved higher AUROC compared to all other methods in most of the cases.

The current practice of Drug Synergy predictions usually considers the molecular structure [13]. However, some recent research has shown that the similarity of the targets, and cell response could also contribute to whether two drugs would have synergism. Sayed-Rzgar H. and Xiaobo Z. conducted numerical experiments to show that the cell response and network information could be used as an input of the prediction model and outperform using structural information as input [10] This research offers us a new perspective to utilize both chemical and bioactivity profiles of small molecules could further improve the Drug Synergy predictions. However, there is still a need for a method to integrate those chemical and bioactivity profile information into one model. In this work, we provide an architecture to compact that information to improve the molecule property prediction. This model has the potential to deploy for the Drug Synergy predictions after modification.

## 5 Conclusion

In summary, the integration of bioactivity profiles with chemical structural information is essential for a comprehensive understanding of small molecules. However, the significant challenge lies in dealing with a large proportion of missing experimental validation data when incorporating these profiles into computational models. In our study, we introduced Bio-Mol, an AI model that effectively addresses this issue by employing a pretrain and finetunes strategy to incorporate high proportions of missing bioactivity profiles during small molecule representation learning. Our results demonstrate a notable improvement in predicting molecule properties. Furthermore, we explored the potential of our model in predicting synergistic drug combinations, highlighting its versatility and broader applications in drug discovery.

## 6 Code availability

The source code of Bio-Mol and the scripts used in the study are available online (https://github.com/YIPhiufung1997/Bio-Mol).

## 7 Acknowledgements

## Notes

### Competing Interest Statement

The authors have declared no competing interest.

## References

1. Bertoni, M., Duran-Frigola, M., i Mompel, P.B., Pauls, E., Orozco-Ruiz, M., Guitart-Pla, O., Alcalde, V., Diaz, V.M., Berenguer-Llergo, A., Brun-Heath, I., Villegas, N., de Herreros, A.G., Aloy, P.: Bioactivity descriptors for uncharacterized chemical compounds. Nature Communications 12(1) (Jun 2021). https://doi.org/10.1038/s41467-021-24150-4, 10.1038/s41467-021-24150-4

2. Delmas, M., Filangi, O., Paulhe, N., Vinson, F., Duperier, C., Garrier, W., Saunier, P.E., Pitarch, Y., Jourdan, F., Giacomoni, F., Frainay, C.: FO-RUM: building a Knowledge Graph from public databases and scientific literature to extract associations between chemicals and diseases. Bioinformatics 37(21), 3896–3904 (09 2021). https://doi.org/10.1093/bioinformatics/btab627, 10.1093/bioinformatics/btab627

3. Dong, Y., Peng, C.Y.J.: Principled missing data methods for researchers. SpringerPlus 2(1) (May 2013). https://doi.org/10.1186/2193-1801-2-222, 10.1186/2193-1801-2-222

4. Dowden, H., Munro, J.: Trends in clinical success rates and therapeutic focus. Nature Reviews Drug Discovery 18(7), 495–496 (May 2019). https://doi.org/10.1038/d41573-019-00074-z, 10.1038/d41573-019-00074-z

5. Duran-Frigola, M., Pauls, E., Guitart-Pla, O., Bertoni, M., Alcalde, V., Amat, D., Juan-Blanco, T., Aloy, P.: Extending the small-molecule similarity principle to all levels of biology with the chemical checker. Nature Biotechnology 38(9), 1087–1096 (May 2020). https://doi.org/10.1038/s41587-020-0502-7, 10.1038/s41587-020-0502-7

6. Fang, X., Liu, L., Lei, J., He, D., Zhang, S., Zhou, J., Wang, F., Wu, H., Wang, H.: Geometry-enhanced molecular representation learning for property prediction. Nature Machine Intelligence 4(2), 127–134 (Feb 2022). https://doi.org/10.1038/s42256-021-00438-4, 10.1038/s42256-021-00438-4

7. Fang, Y., Zhang, Q., Zhang, N., Chen, Z., Zhuang, X., Shao, X., Fan, X., Chen, H.: Knowledge graph-enhanced molecular contrastive learning with functional prompt. Nature Machine Intelligence 5(5), 542–553 (May 2023). https://doi.org/10.1038/s42256-023-00654-0, 10.1038/s42256-023-00654-0

8. Haghighi, M., Caicedo, J.C., Cimini, B.A., Carpenter, A.E., Singh, S.: Highdimensional gene expression and morphology profiles of cells across 28, 000 genetic and chemical perturbations. Nature Methods 19(12), 1550–1557 (Nov 2022). https://doi.org/10.1038/s41592-022-01667-0, 10.1038/s41592-022-01667-0

9. Harrison, R.K.: Phase II and phase III failures: 2013–2015. Nature Reviews Drug Discovery 15(12), 817–818 (Nov 2016). https://doi.org/10.1038/nrd.2016.184, 10.1038/nrd.2016.184

10. Hosseini, S.R., Zhou, X.: CCSynergy: an integrative deep-learning framework enabling context-aware prediction of anti-cancer drug synergy. Briefings in Bioinformatics 24(1), bbac588.(12 2022). https://doi.org/10.1093/bib/bbac588, 10.1093/bib/bbac588

11. Jakobsen, J.C., Gluud, C., Wetterslev, J., Winkel, P.: When and how should multiple imputation be used for handling missing data in randomised clinical trials – a practical guide with flowcharts. BMC Medical Research Methodology 17(1) (Dec 2017). https://doi.org/10.1186/s12874-017-0442-1, 10.1186/s12874-017-0442-1

12. Liu, S., Wang, H., Liu, W., Lasenby, J., Guo, H., Tang, J.: Pretraining molecular graph representation with 3d geometry (2021). 10.48550/ARXIV.2110.07728, https://arxiv.org/abs/2110.07728

13. Preuer, K., Lewis, R.P.I., Hochreiter, S., Bender, A., Bulusu, K.C., Klambauer, G.: DeepSynergy: predicting anti-cancer drug synergy with Deep Learning. Bioinformatics 34(9), 1538– 1546 (12 2017). https://doi.org/10.1093/bioinformatics/btx806, 10.1093/bioinformatics/btx806

14. Rong, Y., Bian, Y., Xu, T., Xie, W., Wei, Y., Huang, W., Huang, J.: Self-supervised graph transformer on large-scale molecular data (2020). 10.48550/ARXIV.2007.02835, https://arxiv.org/abs/2007.02835

15. Sadybekov, A.V., Katritch, V.: Computational approaches streamlining drug discovery. Nature 616(7958), 673–685 (Apr 2023). https://doi.org/10.1038/s41586-023-05905-z, 10.1038/s41586-023-05905-z

16. Takebe, T., Imai, R., Ono, S.: The current status of drug discovery and development as originated in united states academia: The influence of industrial and academic collaboration on drug discovery and development. Clinical and Translational Science 11(6), 597–606 (2018). 10.1111/cts.12577, https://ascpt.onlinelibrary.wiley.com/doi/abs/10.1111/cts.12577

17. Wu, Z., Ramsundar, B., Feinberg, E.N., Gomes, J., Geniesse, C., Pappu, A.S., Leswing, K., Pande, V.: MoleculeNet: a benchmark for molecular machine learning. Chemical Science 9(2), 513–530 (2018). https://doi.org/10.1039/c7sc02664a, 10.1039/c7sc02664a

